# Antibiotic resistance profiles and population structure of disease-associated *Staphylococcus aureus* infecting patients in Fort Portal Regional Referral Hospital, Western Uganda

**DOI:** 10.1101/2020.11.20.371203

**Authors:** G. Ackers-Johnson, D. Kibombo, B. Kusiima, M.L. Nsubuga, E. Kigozi, H.M. Kajumbula, D.P. Kateete, R. Walwema, H.L. Ackers, I.B. Goodhead, R.J. Birtles, C.E. James

**Author notes:** **Corresponding Author:** Chloë E James.

## Abstract

Tackling antimicrobial resistance (AMR) is particularly challenging in low-resource settings such as Fort Portal Regional Referral Hospital (FPRRH) in Western Uganda. Specific knowledge of local AMR epidemiology is required to inform evidence-based improvement of antibiotic stewardship measures in the hospital. To address this, we combined existing antimicrobial susceptibility testing (AST) from FPRRH, with whole genome sequencing (WGS) of 41 *Staphylococcus aureus* isolates (2017-2019). AST revealed 73% (30/41) of isolates were resistant to one or more antibiotics and 29% (12/41) were multi-drug resistant (MDR). Resistance phenotypes were largely explained by the presence of antibiotic resistance genes in WGS data. Five isolates were methicillin-resistant *S. aureus* (MRSA) and MDR. Although all isolates were susceptible to clindamycin, a 24% carriage of *erm* genes suggests potential for rapid development of resistance. We inferred a population structure for the *S. aureus* isolates by comparing their core genomes. Twenty isolates formed a tight cluster corresponding to multilocus sequence typing clonal complex (CC) 152, a CC found to be particularly prevalent in northern Africa. The frequency of genes associated with methicillin, chloramphenicol and ciprofloxacin resistance were significantly lower among CC152 strains than non-CC152 strains; thus, in keeping with previous work, we find that CC152 is almost exclusively methicillin-sensitive *S. aureus* (MSSA). Also, in agreement with other studies, we observed that the occurrence of Panton-Valentine leukocidin toxin-encoding genes was significantly higher among CC152 strains than non-CC152 strains. However, we also observed that the coagulase gene was over-represented in this CC, further defining the virulence strategy of this important pathogen. By generating detailed information about the epidemiology of circulating *S. aureus* and their antibiotic susceptibility, our study has provided, for the first time, data on which evidence-based infection and AMR interventions at FPRRH can be based.

## Introduction

The burden of antimicrobial resistance (AMR) is most keenly felt in low resource settings, where infection control measures are limited and access to antibiotics is intermittent in terms of both availability and type [1]. In Uganda, bacterial infections are responsible for >37% of hospital admissions and despite major international investments in healthcare systems, 90-161 deaths per 100,000 population are caused by infectious disease compared to <38 per 100,000 across the developed world [2]. Inappropriate prescribing, dispensing and administration of antibiotics are likely to contribute to this excess by compromising their utility and increasing the risk of adverse drug reactions and antibiotic resistance [2–6].

Study on the emergence and epidemiology of AMR in *S. aureus* and spread of methicillin resistant *S. aureus* (MRSA) in Sub Saharan Africa (SSA) has intensified [7, 8] but data from resource-poor regions remain sparse. In Uganda, AMR phenotyping of *S. aureus* strains derived from symptomatic and apparently healthy humans and livestock has suggested that resistance to different and multiple antibiotics, including methicillin is prevalent [9–11].

The Ugandan National Action Plan (UNAP) 2018 was developed to address the high prevalence of AMR infections that restrict treatment options across the country. Better surveillance of AMR is a specific objective of the UNAP to strengthen the evidence base for policies on optimal use of antibiotics to preserve their efficacy [6]. The 2017 Joint External Evaluation noted that 25 healthcare facilities are now regularly performing antimicrobial susceptibility testing (AST) in Uganda [12]. However, as these facilities primarily lie in the more heavily populated areas of the country, most AMR surveillance has focused on them (e.g., Kampala [13, 14], Mbarara [15] and Gulu [16]) rather than the more remote, rural settings in which, according to World Bank figures, 76% of the population live [17]. A 2018 study identified spread of antibiotic-resistant *Escherichia coli* bacteria between people, livestock in rural Western Uganda, highlighting the need for further study [18].

To assist capacity in laboratory testing around the UNAP, the Ugandan Ministry of Health and Infectious Diseases Institute (IDI) have recently equipped the Fort Portal Regional Referral Hospital (FPRRH) with a microbiology laboratory capable of routine bacterial identification and AST. FPRRH serves the rural region of Kabarole, in the West of Uganda, near the border with the Democratic Republic of Congo, which is one such area where little data on AMR have been published.

Investigation into the genetic basis of antibiotic resistance and the molecular epidemiology of antibiotic resistant bacteria is needed to gain a deeper understanding of the drivers of resistance spread. PCR-based assays have been used to explore of presence/absence of AMR- and virulence-associated genes among Ugandan *S. aureus* isolates [9, 13, 19], while *spa* (and SCCmec for MRSA) typing has been used to provide an epidemiological context [9, 10, 13, 19–21]. These tools, however, have limited sensitivity; meanwhile advancements in rapid and affordable DNA sequencing technologies, and concurrent improvements of tools to predict AMR *in silico*, have led to improvements in diagnostic microbiology and studies of AMR [22].

This study is part of a multi-disciplinary intervention at FPRRH linking laboratory testing with training of health professionals in understanding common drivers of AMR aiming to reduce unnecessary and ineffective use of antibiotics in line with the UNAP [23]. We combine existing hospital AST data with whole genome sequencing to investigate antibiotic resistance profiles, population structure and virulence gene carriage of *S. aureus* isolated from a range of infection sites including wounds, blood, and the urinary tract, in a region with little previous published data, towards better informed treatment practice and infection/AMR control policies.

## Materials and Methods

### Bacterial isolation and identification

*S. aureus* isolates were routinely collected and phenotypically characterised by the FPRRH microbiology laboratory between 2017 and 2019. These isolates were primarily derived from skin wounds or suspected urinary tract or bloodstream infections. Bacteria were isolated and sub-cultured onto either blood agar, MacConkey agar and chocolate agar (Oxoid) at 37 °C (24-48 h). Putative *S. aureus* isolates were confirmed by Gram staining followed by a positive catalase and Staphaurex latex agglutination test (Thermo Fisher Scientific). All isolates were stored at −80°C. Sample metadata are available in Supplementary Data 1.

### Antibiotic Susceptibility Testing

For each isolate, a single colony was isolated and sub-cultured, prior to screening by a disk diffusion assay according to EUCAST guidelines [24]. Susceptibility to eight different antibiotics belonging to different classes was tested: cefoxitin (30μg), a second-generation cephalosporin used to indicate MRSA; ciprofloxacin (5μg), a fluoroquinolone; cotrimoxazole (sulfamethoxazole - trimethoprim (1.2μg and 23.75μg respectively)); tetracycline (30μg); erythromycin (15μg), a macrolide and clindamycin (2μg), a lincosamide; gentamycin (10μg), an aminoglycoside; and chloramphenicol (30μg) (Oxoid).

### DNA Extraction and Quality Control

Bacterial cultures were grown in Luria Bertani (LB) broth for 18-24h at 37 °C and DNA was extracted using the Isolate II Genomic DNA Kit (Bioline, UK) according to the manufacturer’s instructions, including the extra lysozyme pre-treatment step for efficient lysis of Gram-positive cells. DNA concentration and purity were measured using the Qubit 3.0 Fluorometer (Thermo Fisher Scientific) with Quant-iT dsDNA Assay Kit, High Sensitivity (Invitrogen). DNA library quality control was performed using a Tapestation 2200 with a High Sensitivity D1000 kit (Agilent Technologies) according to the manufacturer’s instructions.

### Whole Genome Sequencing, Draft Genome Assembly and Analysis

Whole genome DNA libraries were prepared using the Nextera XT library preparation kit according to the manufacturer’s instructions. Samples were multiplexed on a single MiSeq flowcell using Nextera XT barcodes and normalised using the bead-based normalisation kit (Illumina). Sequencing of 41 pooled genomes was performed simultaneously at Makerere University, Uganda and the University of Salford, UK, as part of a training activity to support UNAP AMR initiatives. In both cases, DNA sequencing was performed using a MiSeq v2 500 cycle kit (2×250bp reads; Illumina). Raw reads were trimmed using fastq-mcf and assembled using SPADES v3.9.1 [25]. The resulting draft assemblies were annotated using PROKKA v 1.13.3 [26].

The online webtool PathogenWatch (https://pathogen.watch) was used to cluster the annotated genomes of each isolate to predict antibiotic resistance profiles. PathogenWatch uses PAARSNP, bespoke software that uses BLAST against a custom database (combined from public AMR databases such as CARD, ResFinder and NCBI) to highlight the presence or absence of genes and SNPs implicated in antimicrobial resistance and infers their combined effect. Chloramphenicol resistance was predicted by CARD (*Staphylococcus intermedius* chloramphenicol acetyltransferase) analysis [27] implemented in ABRICATE (https://github.com/tseemann/abricate). Virulence gene presence was also predicted using ABRICATE, against the VFDB database [28]. Multi-locus sequence types (MLST) were confirmed against the pubMLST database [29] using the software “MLST” (https://github.com/tseemann/mlst). Core genome alignment was ascertained by SNIPPY [30], and a phylogenetic tree generated by FastTree using a GTR+CAT model of nucleotide evolution [31] (Figure 1). Additional genomic analyses were performed using ABRICATE and ROARY [32].

**Figure 1:**
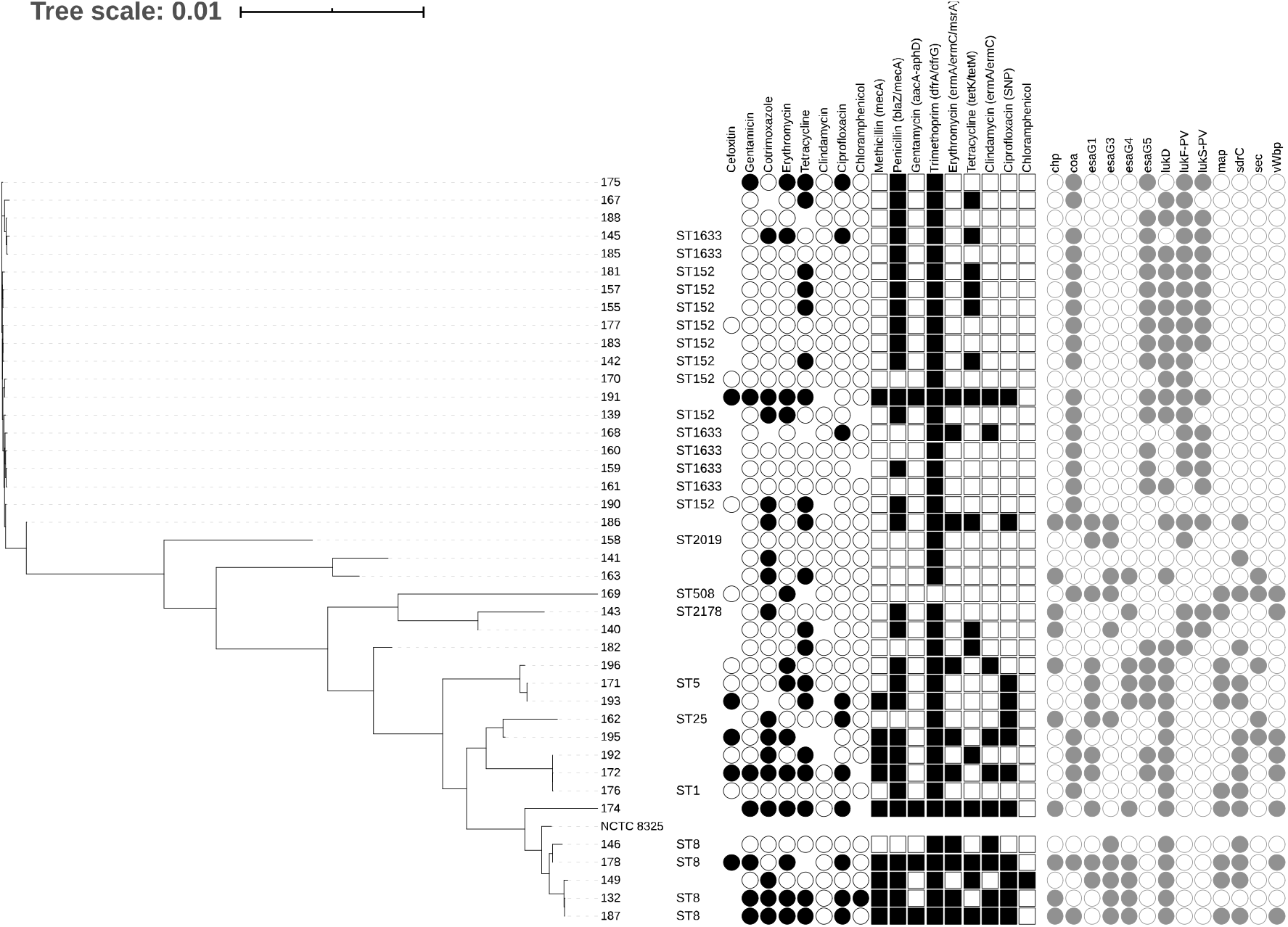
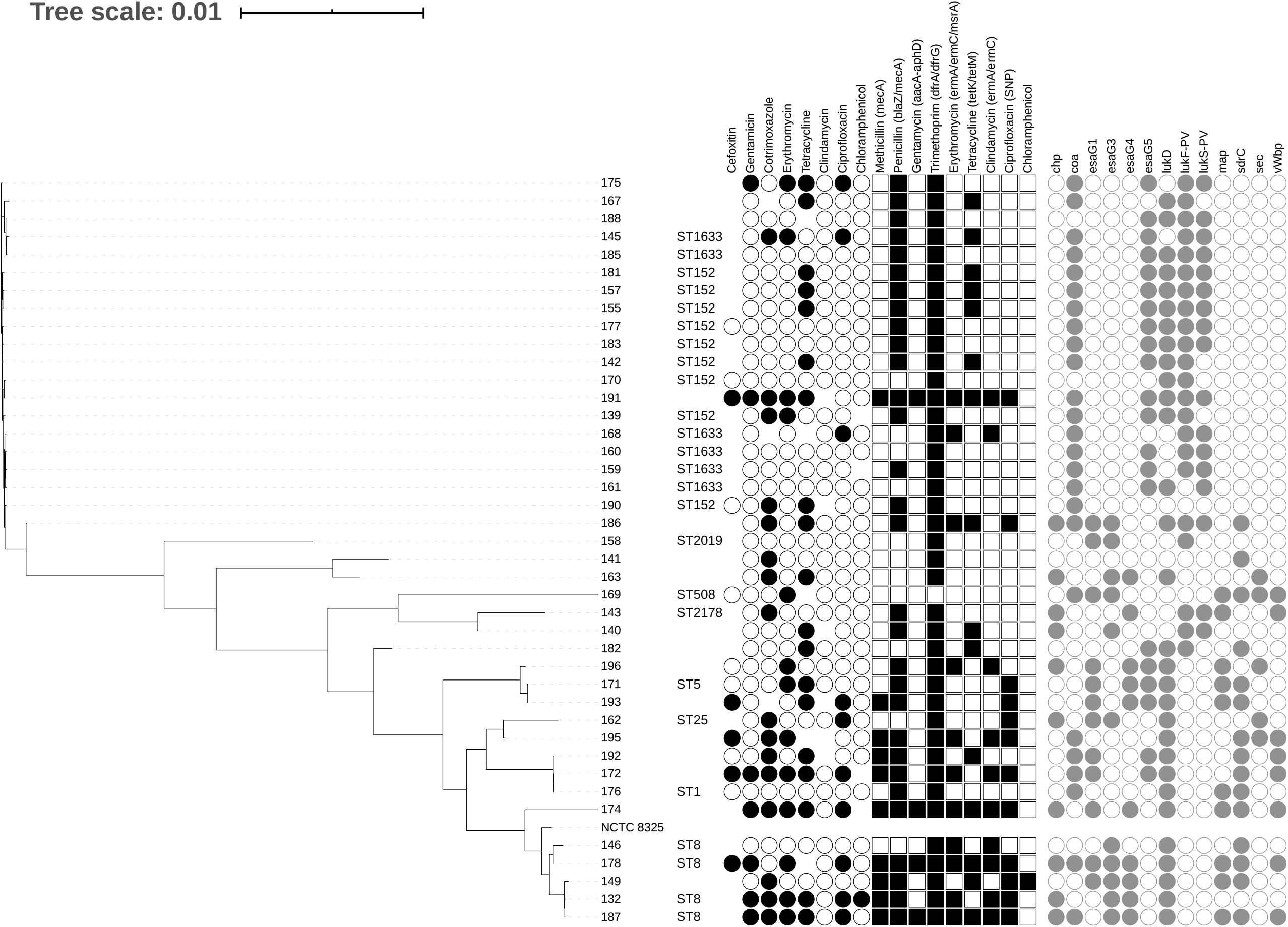
Core genome phylogeny of 41 *Staphylococcus aureus* isolates from Fort Portal Regional Referral hospital isolated between 2017 and 2019, aligned against the NCTC 8325 reference genome [63]. Tree is displayed alongside antibiotic sensitivity data (coloured black circles represent presence, empty circles represent absence, no circle represents no data available); Genotypic antibiotic resistance predictions by PathogenWatch, (coloured squares represent presence of AMR SNP (Ciprofloxacin) or genes (others). For AMR genes shapes represent presence of antibiotic genes / SNP, according to class. The core genome was generated by SNIPPY [30], and the tree file generated using FastTree [31] using a generalized time-reversible (GTR+CAT) model of nucleotide evolution. Also shown are sample ID, Sequence Type (predicted by pubMLST [29]). Virulence gene presence (VFDB) is displayed as grey circles. Further sample metadata can be found in Supplementary Data 1.

Univariant statistical analysis of antimicrobial resistance and virulence gene presence/absence was performed in Minitab.

## Results & Discussion

### Antibiotic susceptibility testing (AST) at FPRRH

This study reviewed the antibiotic resistance profiles of 41 *S. aureus* isolates determined by disk diffusion assay as part of the routine laboratory diagnostic service at FPRRH, Uganda (Figure 1). Overall, 73% of *S. aureus* isolates (30/41) were resistant to one or more antibiotics. 32% (13/41) were resistant to one antibiotic, most commonly tetracycline, and a further 41% (17/41) were resistant to two or more antibiotics, most commonly including cotrimoxazole and tetracycline. 24% of isolates (10/41) were classified as multi-drug resistant (MDR), exhibiting resistance to antibiotics belonging to three or more classes [33]. 5% (2/41) were resistant to antibiotics belonging to six different classes, and thus considered pan resistant. These levels of resistance and MDR are not unexpected; for example, Seni and colleagues [9] reported 21% of Gram-positive isolates from surgical site wounds of patients in Mulago Hospital, Kampala (Uganda) were MDR, whereas Kateete and colleagues [34] observed 31% of *S. aureus* isolates recovered from nasopharyx of children in rural Eastern Uganda were MDR. A study of post-operative sepsis cases in rural Eastern Uganda [35] reported 14% *S. aureus* isolates to be resistant to five different classes of antibiotics and 3% resistant to six different classes [35]. Whilst the Ugandan national action plan (UNAP) reports prevalence of MRSA in healthcare settings at 2-50% [23], the WHO AMR global report on surveillance (2014) found overall MRSA prevalence to range from 12-80% across Africa and to exceed 20% in all six WHO regions [36]. There have been reports of more than 80% MRSA prevalence in African, American and Western Pacific regions [36].

Empirical treatment (ceftriaxone + metronidazole) is prescribed for suspected infections at FPRRH. Whilst metronidazole covers anaerobic bacteria, ceftriaxone is a broad-spectrum cephalosporin that is well tolerated and commonly used to effectively treat infections caused by most aerobes including methicillin susceptible *S. aureus* (MSSA). Cefoxitin susceptibility testing was used as a proxy to indicate resistance to ceftriaxone and identify MRSA. However, limited resources at FPRRH prevented a considerable number of isolates from being tested. Of 13 isolates tested, five (38%) were resistant (MRSA), a prevalence akin to previous Ugandan studies [9, 16, 35]. All five MRSA isolates were MDR; Kateete and colleagues [13] also observed markedly higher rates of MDR among MRSA strains compared to MSSA strains. The observed frequency of MRSA at FPRRH suggest a potentially high level of failure of the empirical treatment, and the need for informed decisions about procurement of alternative antibiotics.

### Whole genome sequencing data

Draft whole genomes were generated for all 41 *S. aureus* isolates using Illumina MiSeq data. All sequence read data used for genome assembly, annotation and AMR genotype prediction are available on the European Nucleotide Archive under project accession: PRJEB40863. According to PathogenWatch (data available at [37]), we identified between 86% to 100 % of the core gene families for all isolates (mean 95% per isolate). N50 ranged from 3,045 bp to 145,228 bp (mean 26.8 Kbp; See Supplementary Data 1).

### Genomic correlation with phenotypic AMR profiles

Although the primary aim of this study was to predict antibiotic resistance data for *S. aureus* isolates for which phenotype data was lacking due to poor resources, the study provides an opportunity to correlate AMR phenotype with genotype prediction. A recent study of 778 MRSA isolates from the UK, comparing Illumina WGS data – based analysis with disc diffusion AST data showed 99.7% accuracy [38]. To that end, we have compared AST results from FPRRH with antibiotic resistance gene (ARG) presence/absence data drawn from WGS. Genetic markers of resistance were found in all 11 (27%) isolates that were reported as susceptible to all the antibiotics tested for at FPRRH. Phenotypic and genotypic predictions of susceptibility to each antibiotic were compared in turn (Figure 1 and Supplementary Data 1).

Tetracycline resistance was the most common phenotype found in 56% of isolates tested (19/34). Previously, prevalence of tetracycline resistance in Uganda has been reported as high as 55% among clinical isolates [39] and 68% among environmental/carriage isolates [13]. Tetracycline resistance was accounted for in 67% (12/18) of genomes by the presence of *tetK/M* (encoding an efflux pump and ribosomal protection respectively). However, no tet genes were found in the genomes of seven tetracycline resistant isolates. Tetracycline susceptibility was predicted (by absence of *tetK/M* genes) for 93% of the susceptible isolates tested (14/15). Tetracycline resistance correlation between WGS and AST data has been reported as high as 100% [38]. Others report 83% correlation of resistance phenotypes with presence of *tetK* and *tetM* genes [40]. Although tetracycline dispensing for these infection types has not been reported at FPRRH, it is commonly used in treatment of Ugandan livestock, as a broad-spectrum antibiotic, antiparasitic and as a growth supplement [41]. Nevertheless, resistance levels in livestock in western Uganda have been reported as low as 8% [42] or 12% [41], implying possible widespread but unrecognised dispensing of this antibiotic to humans [43].

Of 37 isolates tested 16 (43%) were resistant to cotrimoxazole (trimethoprim + sulfamethoxazole) and a further 5 (14%) showed an intermediate result. However, the remaining 16 (43%) were susceptible despite genetic indicators of trimethoprim resistance (*dfrG/dfrA*) present in almost all isolates (98%, 40/41). Others have reported trimethoprim resistant phenotypes where isolates were genotypically predicted to be susceptible, in one case despite the presence of a *dfrA* gene, suggesting that *in silico* prediction of trimethoprim resistance may require further work [38, 44].

Increasing levels of *dfrG*-related trimethoprim resistance were reported in 2014, suggesting that resistance would soon be widespread in Africa through imported infections [45]; This study would suggest that in the prevailing six years, this prediction has come to pass, but does not reflect the much higher resistance rates reported elsewhere in Uganda [8, 9, 13, 16, 46]. This may have been driven, in part, by the widespread use of cotrimoxazole for prophylaxis against secondary infections in HIV positive patients across SSA [47, 48] and highlights the need for wider study and control of other antibiotics showing decreased efficacy.

AST suggested that 71% of isolates (29/41) were susceptible to erythromycin and these phenotypes matched the genotype in most cases (26/29). Thirteen isolates (32%) were found to be resistant to erythromycin, which is considerably lower than reported by several other Ugandan studies [35, 46, 49, 50] and all erythromycin-resistant isolates were susceptible to clindamycin. For some isolates, this could be explained by the presence of an inducible efflux pump gene (*msrA*), which is implicated in resistance to erythromycin [51]. More commonly, resistance to both these antibiotic groups is partly predicted by the presence of *erm* genes, the products of which modify the ribosomal target site to prevent antibiotic binding. Erythromycin resistance (predicted by PathogenWatch by detection of *ermA* (1/41), *ermC* (11/41) or *msrA* (5/41) genes) did not always correlate with phenotype: 38% (5/13) of resistant isolates were not predicted by genotype. In contrast, 100% of isolates tested (35/35) appeared susceptible to clindamycin despite detection of *ermA* or *ermC* in 12 isolates. This suggests that the genes are under erythromycin-inducible regulation, but that exposure to clindamycin *in vivo* could select for constitutive expression mutants, that would confer resistance to both [52]. According to the WHO essential medicines list (2019), clindamycin is an access antibiotic recommended as a second choice for bone / joint infections. Clindamycin has not been used at FPRRH but has been used elsewhere in Uganda; a 2014 study based at Mbarara Regional Referral Hospital in Southwestern Uganda reported 36% of isolates as clindamycin-resistant and also showed discrepancies between phenotypes and genotypic prediction [49]. Clindamycin resistance rates of 0-59% have been reported elsewhere in Uganda between 2009-2018 [46] and was reported as early as 2008 in Tanzania [53]. It is therefore clear that erythromycin/clindamycin resistance is a complex issue that requires ongoing monitoring given the potential to expanded resistance.

Other options from the WHO list of access antibiotics include gentamycin and chloramphenicol, which are often recommended when empirical treatments fail [46]. Gentamycin is a commonly used aminoglycoside at FPRRH; 17% of isolates (7/41) were gentamycin resistant, but specific aminoglycoside resistance genes (*apH3* and *aacA-aphD*) were only found in 4 (of 41) genomes. Similar resistance rates have been reported in Kampala [9] and Gulu [16], but others report much higher rates [46].

Aminoglycoside resistance is complex, and conflicts between genotype and phenotype have been previously reported [54], but may also be representative of the short read sequencing, which could be clarified using long-read technologies or complementary techniques. Phenotypic and genotypic analysis of chloramphenicol susceptibility were largely in agreement, with 97% of isolates being susceptible to chloramphenicol (36/37) CARD analysis predicted only one isolate (sample 149) to be resistant, which did not correlate with several other studies from Mulago hospital, Kampala that suggest much higher rates of chloramphenicol resistance ranging between 14% and 88% [9, 13, 14, 46, 50].

Ciprofloxacin (a fluoroquinolone) and ceftazidime (a third-generation cephalosporin) both appear on the WHO watch list of essential antibiotics that are considered at high risk of selecting for further bacterial resistance, and should be prioritised for stewardship programmes [2]. A total of 24% (10/41) of *S. aureus* isolates at FPRRH were resistant to ciprofloxacin, and two more exhibited intermediate resistance; 75% of these correlated with genotypic prediction with mutations in *gyrA* and *grlA* genes. Ciprofloxacin resistance in *S. aureus* is known to develop rapidly through single nucleotide mutations [55], and rates vary across other Ugandan studies, most of which report higher rates of ciprofloxacin resistance than at FPRRH [9, 14, 16, 46].

Phenotypic resistance to cefoxitin, as a proxy for ceftazidime resistance and MRSA corresponded to genotypically-predicted methicillin resistance, with one susceptible isolate being predicted to be resistant. Of note, stocks of cefoxitin disks were poorly maintained at FPRRH during the study period and 28 isolates were not able to be tested. Of 13 isolates tested, five (38%) were resistant (MRSA). In contrast to the rates of resistant isolates to other antibiotics, the percentage of MRSA at FPRRH is similar or higher than those reported elsewhere (Kampala: 37.5% [9], 29% [46] and 28% [39]; Mbale: 27% [35] and Gulu: 22% [16]). All five MRSA isolates from FPRRH were MDR. 14% (4/28) of isolates that were not tested harboured *mecA*, a clear genetic indicator of MRSA. The majority (71%) of isolates (30/41) also harboured the *blaZ* gene which encodes a β-lactamase conferring resistance to a range of penicillins and other β-lactam antibiotics. Two such isolates remained susceptible to ciprofloxacin, which could be recommended as an alternative treatment in these cases.

Overall, the WGS data provided a more detailed view of the AMR mechanisms circulating in *S. aureus*-associated infections at FPRRH although susceptibility was more commonly predicted than resistance. Given the non-contiguous nature of the draft genomes this could be due to a lack of complete gene coverage enabling confident prediction of gene presence or absence. Complete genomes, such as using PacBio or Oxford Nanopore sequencing would increase confidence in prediction, but the additional effort may not be cost-effective in low-middle income resource settings. These data are nevertheless useful for in-filling missing information from AST data due to the poor supply of appropriate materials, towards informing more-effective treatment of *S. aureus* infections at FPRRH.

### Population structure of FPRRH *S. aureus* isolates: domination of MSSA CC152

We observed marked diversity among the 41 *S. aureus* isolates we characterised, as quantified in a core genome phylogeny (Figure 1). We were able to obtain complete MLST data for 25 of these (Figure 1) and delineated nine different STs belonging to six CCs. Isolates belonging to a particular CC clustered together in the population structure inferred from core genome comparison (Figure 1). Most isolates for which we obtained complete MLST data (15) belonged to ST152 and its single locus variant ST1633 (CC152). For three more isolates, we obtained MLST data for 6 out of 7 loci allowing them to be attributed to CC152 if not a specific ST. Two further isolates (180 and 191) had more incomplete MLST data, but core genome analysis indicated they clustered tightly with CC152 isolates. Thus, in total 20 of the 41 isolates we studied (49%) were likely CC152. Ruimy and colleagues [56] were the first to highlight the exceptionally high frequency of CC152 in northern Africa, having assigned 24% of isolates recovered from the nasopharynx of apparently healthy Malian people to this CC. Numerous subsequent studies of clinical isolates across the continent have supported this observation (reviewed in [57]); most recently, Kyany’a et al (2020) and Obasuyi et al (2020) reported CC152 was the most common clonal complex encountered, representing about a quarter of the clinical isolates they characterised in Kenya and Nigeria respectively [58, 59].

A high proportion of CC152 isolates had either no resistance phenotype or were resistant to just one antibiotic (most commonly tetracycline). 25% were resistant to cotrimoxazole and 20% were resistant to erythromycin. The most common ARGs in this group were *dfrA/dfrG* (100%), *blaZ* (80%) and *tet* (40%). Phenotypic and genotypic screening determined low frequencies of resistance to ciprofloxacin, clindamycin and chloramphenicol. Comparison of CC152 and non-CC152 isolates in this study revealed PathogenWatch predictions of methicillin, chloramphenicol and ciprofloxaxin resistance were all significantly less prevalent in CC152 members (χ^2^ =7.96, 4.39 and 7.00, P=0.005, 0.036 and 0.008 respectively).

PathogenWatch also predicted that resistance-associated *mecA* (methicillin resistance), *grlA* and *gyrA* (ciprofloxacin resistance) were significantly less prevalent among CC152 members (χ^2^ =7.96, 5.42 and 5.63, P=0.005, 0.020 and 0.018 respectively). Thus 19/20 (95%) of CC152 isolates were MSSA. In contrast to other CC152 members, three isolates exhibited MDR phenotypes. Isolates 145, 175 and 191 were resistant to 3, 4 and 5 classes of antibiotics respectively, however, there was little similarity in their resistance profiles and ARG compliments, suggesting that they were acquired independently of one another. Isolate 191 was the only MRSA and harboured genetic markers of resistance to eight different classes of antibiotic. All three MDR isolates were resistant to erythromycin, but associated genetic markers were only present in 191, harbouring both *erm* and *msrA* genes that are commonly plasmid associated. Further investigation is needed to determine how these MDR and potentially pan-resistant profiles have developed in a clonal complex that is mainly associated with healthy individuals.

### FPRRH *S. aureus* CC152 isolates possess a specific virulence gene repertoire

Given the frequency with which CC152 strains were encountered, we explored variation in virulence-associated gene content between isolates belonging to this clonal complex (n = 20) and those that did not (n = 21). The majority of virulence-associated genes were neither over-nor under-represented among CC152 stains, but there were a handful of exceptions. Genes that were significantly more common among CC152 strains included *lukF-PV* and *lukR-PV* that encode the two subunits of the Panton–Valentine leukocidin (PVL) cytotoxin (χ^2^ =20.74, P <0.001 and χ^2^ =18.09, P<0.001 respectively) and *coa*, that encodes a coagulase (χ^2^ = 13.82, P<0.001). An association between CC152 and PVL has been noted previously [56]. Coa is a long-established virulence factor of *S. aureus* that is able to induce blood coagulation by interacting with fibrinogen and prothrombin [60]. Interestingly, von Willebrand factor binding protein, encoded by *vWbp*, exhibits multiple similarities in terms of coagulation induction function to Coa [60]. That *vWbp* is absent from CC152 isolates, but present in about 37% of non-CC152 isolates (χ^2^ =9.47, P=0.002) may point to the two proteins fulfilling similar roles in different *S. aureus* lineages. Several other virulence factor-encoding genes were encountered significantly less frequently among CC152 isolates than non-CC152 isolates. These included genes encoding (i) the chemotaxis inhibitory protein Chp (χ^2^ =7.96, P=0.005), (ii) the adhesin Map (χ^2^=12.60, P=0.003), the serine-aspartate repeat protein SdrC that is associated with biofilm formation (χ^2^=16.79, P<0.001), and the Sec enterotoxin (χ^2^= 5.42, P=0.020). The EsaG protein is thought to serve as an anti-toxin in the *Ess* Type 7 secretion system of *S. aureus* [61]. Multiple genes encoding EsaG have been encountered in *S. aureus* genomes and we observed significant variation in their presence/absence among CC152 members and other isolates, such that some copies (*esaG1*, *esaG3*, and *esaG4*) were under-represented (χ^2^= 11.11, 9.48, 12.60, P <0.001, 0.002, <0.001 respectively) among CC152 strains whereas others (*esaG5*, *esaG7* and *esaG8*) were over-represented (χ^2^= 8.84, 5.03, 6.55, P= 0.003, 0.025, 0.010 respectively). The significance of this is unclear as the role of different copies is unknown.

## Conclusions

The main goal of this study, in keeping with the UNAP, was to create local knowledge of circulating AMR in disease-associated *S. aureus*, to strengthen the evidence base for policies on optimal use of antibiotics to preserve their efficacy, and through which we could inform treatment policy at FPRRH and its surrounding catchment area. Key findings include the incidence of MDR and pan-resistant isolates, the prevalence of PVL and coagulase-carrying CC152 strains in a clinical setting, and the apparent low frequency of resistance to clindamycin and chloramphenicol compared to other regions of Uganda.

The resistance profiles observed in this study were largely similar to those described elsewhere in Uganda. Nevertheless, the largest discrepancies between this and other studies were the universal susceptibility to clindamycin at FPRRH, whilst a relatively high 41% resistance was found at Mulago, and for trimethoprim (43% in this study, and ≥89% in Mulago), and chloramphenicol, for which although rates of resistance vary between 14% - 88% elsewhere [9, 13, 14, 46, 50], FPRRH strains were almost all susceptible. National advice on control strategy based on national clinical guidelines, or data that may be focused upon major urban centres, are not always aligned with antibiotic susceptibility observed at the local level, emphasising the requirement for local knowledge such as that described here [16].

It is clear that procurement of sufficient materials to perform AST tests for resistances identified in the antibiogram, supported by molecular data, will be important for robust future surveillance to inform effective treatment and to ensure a rapid, appropriate planned response to treatment failures. Given the frequent circulation of *S. aureus* isolates between community and hospital settings [62], and combined with the variability in resistance rates across Ugandan studies this further highlights the need for continued, robust epidemiology at the local level to inform policy on control strategies that can minimise treatment failures and the spread of antimicrobial resistance.

## Acknowledgements

This work was supported by Knowledge for Change (registered charity no. 1146911), The University of Salford, Santander and a THET Commonwealth Partnerships for Antimicrobial Stewardship award to HLA (AMSB03). We are grateful to staff and services at Fort Portal Regional Referral Hospital and Makerere University, with particular thanks to Dr Robert Ssekitoleko for meeting coordination in Kampala.

